# Extracellular Matrix Remodeling Associated with Bleomycin-Induced Lung Injury Supports Pericyte-To-Myofibroblast Transition

**DOI:** 10.1101/2020.11.16.384776

**Authors:** Riley T. Hannan, Andrew E. Miller, Ruei-Chun Hung, Catherine Sano, Shayn M. Peirce, Thomas H. Barker

**Author notes:** **Corresponding Author:** Thomas H. Barker.

## Abstract

Of the many origins of pulmonary myofibroblasts, microvascular pericytes are a known source. Prior literature has established the ability of pericytes to transition into myofibroblasts, but provide limited insight into molecular cues that drive this process during lung injury repair and fibrosis. Fibronectin and RGD-binding integrins have long been considered pro-fibrotic factors in myofibroblast biology, and here we test the hypothesis that these known myofibroblast cues coordinate pericyte-to-myofibroblast transitions. Specifically, we hypothesized that αvβ3 integrin engagement on fibronectin induces pericyte transition into myofibroblastic phenotypes in the murine bleomycin lung injury model. Myosin Heavy Chain 11 (Myh11)-CreERT2 lineage tracing in transgenic mice allows identification of cells of pericyte origin and provides a robust tool for isolating pericytes from tissues for further evaluation. We used this murine model to track and characterize pericyte behaviors during tissue repair. The majority of Myh11 lineage-positive cells are positive for the pericyte surface markers, PDGFRβ (55%) and CD146 (69%), and display typical pericyte morphology with spatial apposition to microvascular networks. After intratracheal bleomycin treatment of mice, Myh11 lineage-positive cells showed significantly increased contractile and secretory markers, as well as αv integrin expression. According to RNASeq measurements, many disease and tissue-remodeling genesets were upregulated in Myh11 lineage-positive cells in response to bleomycin-induced lung injury. *In vitro*, blocking αvβ3 binding through cyclo-RGDfK prevented expression of the myofibroblastic marker αSMA relative to controls. In response to RGD-containing provisional matrix proteins present in lung injury, pericytes may alter their integrin profile. This altered matrix-integrin axis contributes to pericyte-to-myofibroblastic transition and represents a possible therapeutic target for limiting the myofibroblastic burden in lung fibrosis.

**Highlights:**

- Pericyte lineage model enables study of transdifferentiating pericytes
- High dimensional flow cytometry used to characterize pulmonary stromal cells
- Pulmonary pericytes express matrix-remodeling genes and proteins in lung injury
- Myofibroblasts derived from pericytes have active αvβ3 integrin
- *In vitro* assay reveals necessity of RGD for pericyte transdifferentiation

## Introduction

Acute lung injury most often leads to a transient activation of resident cells, tissue remodeling, and eventual injury resolution. However, under certain circumstances acute injury can progress into pulmonary fibrosis, a disease characterized by scar buildup and concomitant reduction in functional measures of respiration. These pathologies, such as pulmonary fibrosis, have largely unknown etiology and extremely limited palliative therapeutics [1]. Pulmonary fibrosis is specifically characterized by a reduction in vital respiratory metrics and a persistent wound repair environment consisting of inflammatory cytokines, early and late provisional extracellular matrix (ECM) proteins like fibrin, fibronectin and collagens, and ECM-remodeling enzymes in the lung [2–6]. Cellular infiltration, proliferation, and the expansion of interalveolar spaces in early fibrosis is referred to as fibroproliferation, which is the phase of disease wherein quiescent cells become activated and involved in the fibrotic process [7]. Through the exploration of these activated cells, there is the promise of understanding how transitions to a more chronic fibrotic remodeling program may occur.

The historical example of an activated, fibrotic effector cell is the myofibroblast. Myofibroblasts are defined by in situ observation of secretory, contractile, and tissue-remodeling phenotypes, typically through immunohistologic methods. There are no reliable lineage markers for myofibroblasts, as they derive from a variety of quiescent cell populations, the diversity of which can lead to vast differences in regeneration and tissue remodeling outcomes. Thus, recent research into tissue-resident stromal cell populations have focused on identifying and characterizing the various myofibroblast progenitor populations [8–12].

One known myofibroblast progenitor population is the perivascular mural cell, or pericyte, a cell physically associated with microvascular endothelial cells in capillary networks. Pericytes are phenotypically diverse and are typically identified by a variety of surface markers including CD146, PDGFRβ, NG2, and Desmin [13–16]. Pericyte investment in the microvasculature supports vessel integrity and is essential for vascular homeostasis and functional tissue regeneration after insult [14,17]. Pericytes have demonstrated phenotypic plasticity, acting as a source of myofibroblasts in fibrotic disease [18] and other pathologies [19–21]. The myofibroblastic pericyte can emerge in response to lung injury [18,22], responding to classic myofibroblast-promoting conditions, including TGFβ and ECM stiffness [22], two stimuli known to activate classically-defined myofibroblasts. Study of the molecular mechanisms involved in mechanotransduction and activation of latent TGFβ have identified the integrins as essential components in myofibroblastic activation [23].

Integrins are a class of heterodimeric transmembrane receptors which bind to a variety of ligands, the majority of which are found in the ECM. Specific integrin and ligand combinations can potentiate a range of cellular behaviors ranging from differentiation to apoptosis to extravasation. Fibroblast signaling through the αvb3 integrin is thought to be at equilibrium with signaling through α5β1, and when this balance is disrupted in disease (known as an ‘integrin switch’), greater αvb3 integrin signaling drives disease phenotypes [24–26]. It is thought that this shift towards pro-myofibroblastic αvb3 signaling is derived from the increase in Arginine, Glycine, and Aspartate (RGD) ligand found in the fibronectin-rich provisional matrix in early stages of tissue remodeling[25,27–31]

Integrins are no less important in mediating the responses of pericytes to their biochemical and biomechanical environments. The loss of pulmonary basement membrane, in which healthy pericytes are situated, is considered a hallmark of mature and non-resolving fibrosis [32]. Pericyte investment in the basement membrane and capillary network is facilitated by laminin binding to α6 heterodimers, α6β1 and α6β4 [33–35]. For pericytes, the transition from laminin- and collagen IV-rich basement membrane to a fibronectin-rich provisional matrix during early lung injury could invoke a stark change in integrin signaling, similar to the fibroblast integrin switch, leading to phenotypic switching [36]. Indeed, when αv integrin was selectively knocked out via use of a PDGFRβ-cre mouse, its loss was shown to be protective in a bleomycin lung injury model [37].

However, a direct linkage between ECM ligand, surface integrin expression, and pericyte-to-myofibroblast transition has yet to be explored, and whether fibronectin is sufficient to trigger the pericyte-to-myofibroblast transition is an open question. Additionally, characterization of the myofiboblastic pericyte *in vitro* and *in vivo* is typically limited to assessment of a single marker, such as alpha smooth muscle actin (αSMA), limiting our understanding of the broader phenotypic changes that the transitioning pericyte has undergone. Therefore, the goals of this study were to: 1) more comprehensively characterize the phenotypes of pulmonary pericytes and their local ECM environment following lung injury with bleomycin, and 2) test the hypothesis that RGD-mediated integrin signaling can precipitate the pericyte-to-myofibroblast transition.

## Results

### Myhłł lineage reporter mouse labels pericytes in the lung microvasculature

The induction of *Myh11-CreERT2 ROSA STOPfl/fl tdTomato* mice (described in Supplemental Figure 1A) with tamoxifen induces recombination and expression of tdTomato in pericytes, as well as vascular and bronchiolar smooth muscle cells (Figure 1A), consistent with prior work using Myh11 reporter mice [38–41]. The use of the tdTomato fluorescent reporter with the Myh11 Rosa26 construct allows for greater sensitivity in detecting Myh11 lineage-positive cells than the previously published eYFP fluorescent reporter lineage mouse. While observation of Myh11 lineage-positive cells in the pulmonary capillary bed has only been associated with injury in the eYFP reporter mouse [41], we can clearly identify the Myh11 lineage-positive pericytes as being tissue-resident cells before injury. These tdTomato-expressing, fluorescent pericytes become much brighter in disease models, as demonstrated by the differences in relative brightness between saline and bleomycin-treated lungs given the same confocal image acquisition settings in Figure 2A and prior literature [41].

**Figure 1.**
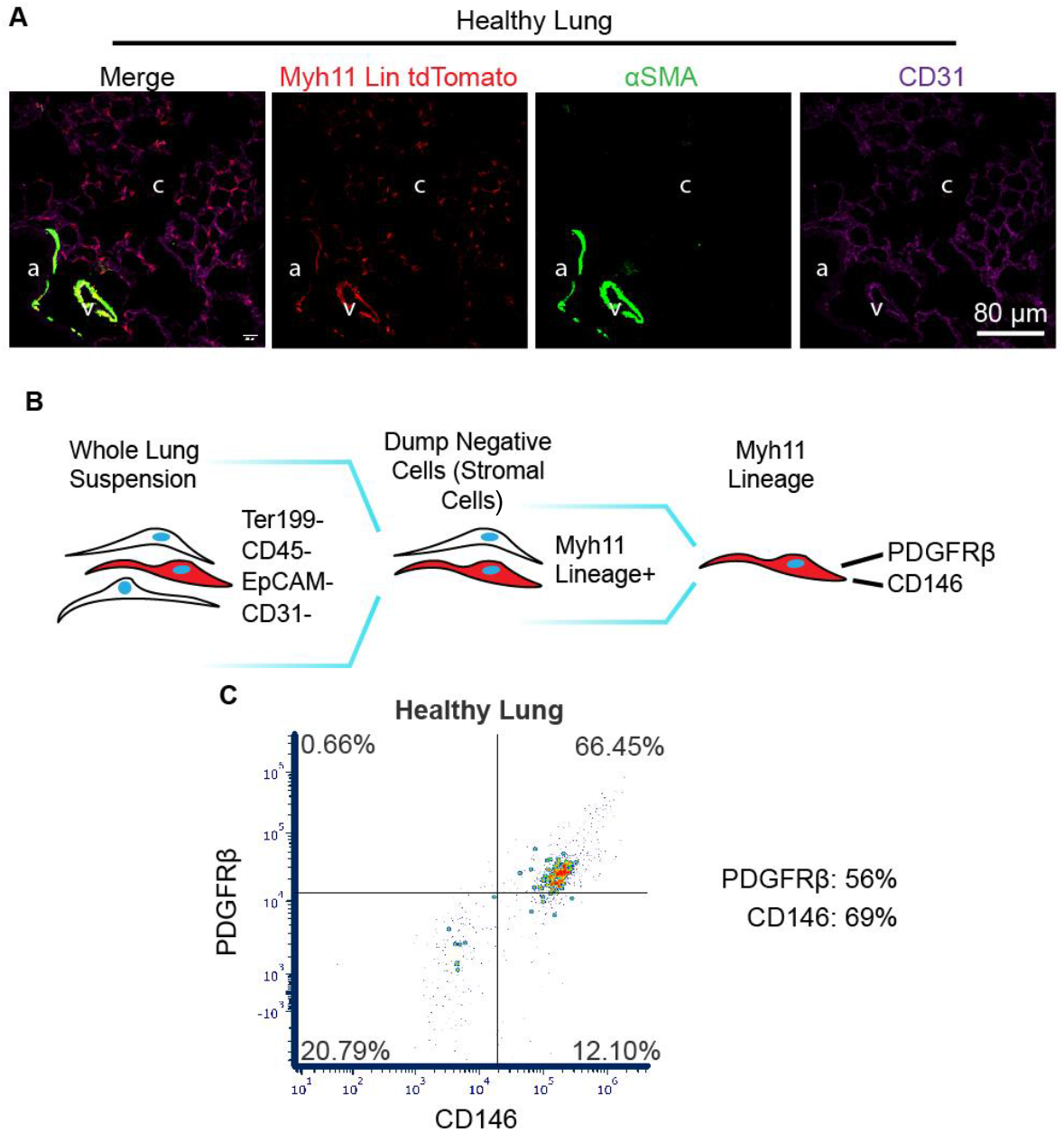
The Myh11-CreERT2 ROSA STOPfl/fl tdTomato reporter mouse labels pericytes in the lung capillary bed. (A) Representative immunofluorescence (IF) micrographs of lung sections stained for tdTomato (red, endogenous fluorophore), αSMA (green), and CD31 (purple). Anatomical structures are denoted “a” for bronchiolar airway lumen, “v” for venule, and “c” for the alveolar capillary bed. (B) Gating hierarchy to isolate Myh11 lineage-positive cells for phenotyping. (C) Representative scatter plot of PDGFRβ and CD146 surface markers on the Myh11 lineage. An average of 55.5% of Myh11 lineage-positive cells in healthy mice were positive for PDGFRβ, while an average of 69.3% were positive for CD146 (n=3).

**Figure 2.**
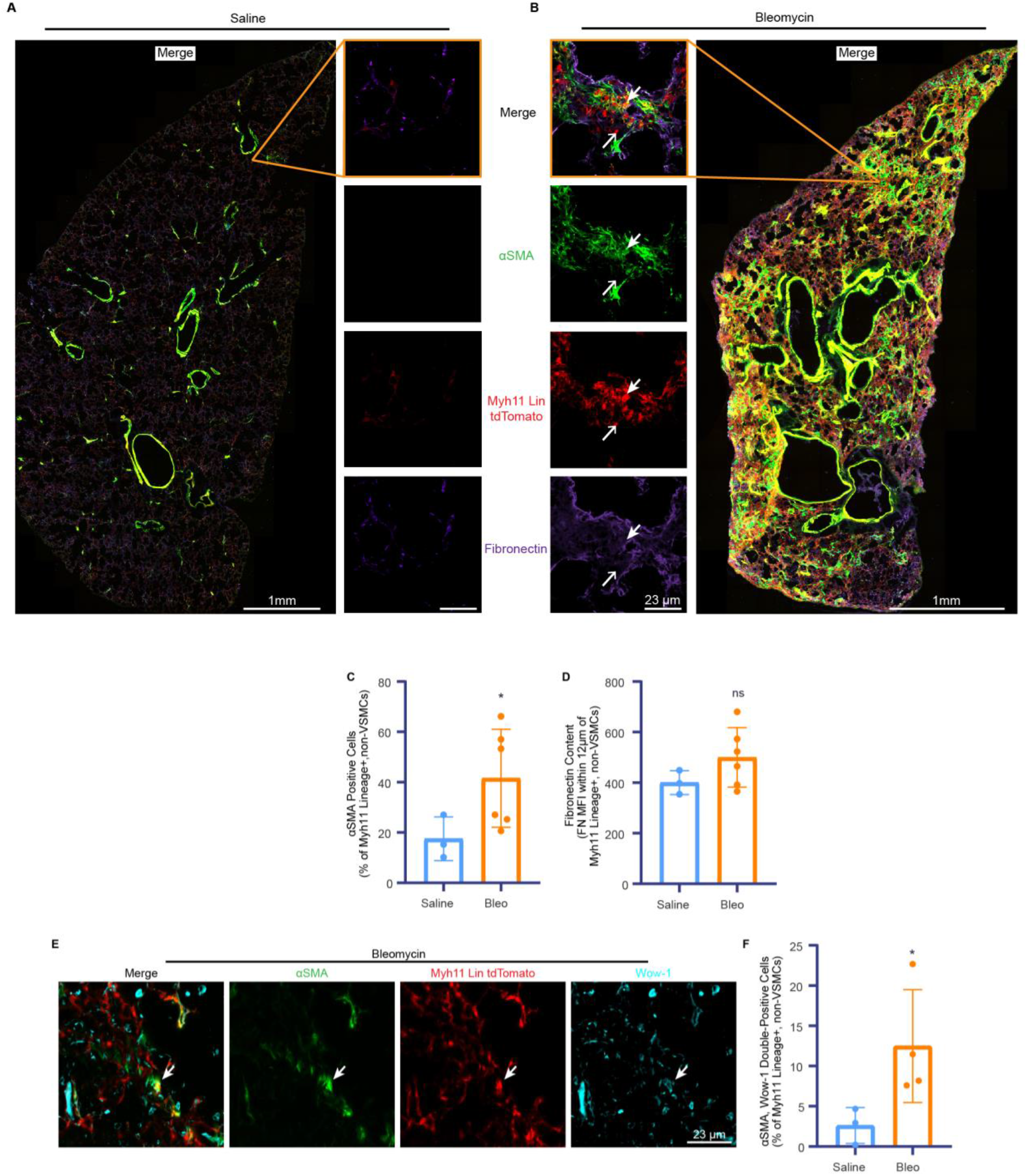
Immunofluorescence (IF) micrographs of lungs from saline and bleomycin treated mice lungs showed increases in perivascular αSMA and engaged αvβ3 integrin. (A,B) Representative confocal micrographs of transverse lung sections immunolabeled for αSMA (green), tdTomato (red, endogenous fluorophore), and fibronectin (purple) from saline (A, n=3) and bleomycin (B, n=6) treated mice. High-magnification inserts (middle) allow for identification and quantification of individual αSMA-positive pericytes (thick arrow) and αSMA-negative pericytes (thin arrow). (C) The number of Myh11 lineage-positive pericytes expressing αSMA is reported as a percentage of the number of total Myh11 lineage-positive pericytes counted across an entire lung section and the mean fluorescence intensity of fibronectin within 13 microns of each pericyte was measured (D). (E) Representative micrograph of Wow-1 staining in the bleomycin treated lung, with an αSMA, Wow-1 double positive pericyte shown (thick arrow) and quantitative comparison between saline (n=3) and bleomycin (n=5) lung sections (F). Data are expressed as means ± standard deviation. Statistical significance was determined via unpaired, one-tailed student’s t-test. ns = not significant; p < 0.05 = *; p < 0.01 = **; p < 0.001 = ***.

Spontaneous recombination is not seen in uninduced mice prior to experimentation (Supplemental Figure 1B). These Myh11 lineage-positive pericytes in the capillary bed extend abluminal processes along capillary endothelium (Figure 1A). The majority location, morphology, surface markers, and body of prior work on this Myh11 lineage [39–42] provide robust evidence to support a classification of Myh11 lineage-positive cells as pericytes.

### Myh11 lineage-positive pericytes adopt myofibroblastic phenotypes within regions of fibroproliferative repair in the injured lung according to immunofluorescent histologic analyses

Using a single-dose intratracheal bleomycin lung injury model, immunofluorescent imaging and analyses were performed on lung specimens from saline-treated control mice and bleomycin-treated mice. Confocal micrographs of transverse sections taken from the midline left lung demonstrate the pronounced tissue remodeling characteristic of the bleomycin disease model (Figure 2A, B), where the interstitial tissue expands through fibroproliferation and ablates the alveolar airspaces [7,43]. This increase in tissue density and loss of alveolar spaces is known to be potentiated by myofibroblastic tissue remodeling. In saline-treated control lungs (Figure 2A), the vast majority of αSMA content can be found in the smooth muscle cells lining larger vessels (pulmonary venules and bronchioles), while more diffuse and non-luminal αSMA is abundant in the bleomycin treated lung (Figure 2B). The proportion of Myh11 lineage-positive pericytes in lung sections expressing αSMA more than doubles two weeks post-bleomycin treatment (Figure 2C). This analysis manually excludes Myh11 lineage-positive vascular smooth muscle cells in bronchioles or venules, as described in the Methods section. An analysis of fibronectin levels local to Myh11 lineage-positive pericytes (within 12 microns of cell soma) revealed no significant difference in fluorescence intensity between saline-treated and bleomycin-treated lungs (Figure 2D). Active perivascular αvβ3 integrin (Figure 2E, Wow-1) increases in bleomycin-treated lung, with αSMA+/Wow-1+ pericytes significantly increasing in frequency in bleomycin-treated lung (Figure 2F).

### Myh11 lineage-positive pericytes isolated from fibrotic lungs show increases in tissue-remodeling markers by flow cytometry

Myh11 lineage-positive pericytes are defined here as live cells negative for the cell-surface markers of other cell lineages Ter119 (erythrocytes), CD45 (myeloid lineage), EpCAM (epithelial cells), CD31 (endothelial cells), which we refer to as “dump negative”, and positive for Myh11 lineage and CD146 (pericyte marker). Cells were isolated from whole-lung digestions from bleomycin-treated and saline-treated lungs, as depicted in Figure 3A. Myh11 lineage-positive pericytes were evaluated for a panel of matrix-remodeling and matrix-binding markers, including: αSMA, Collagen type 1 alpha 1 (Col1a1) and integrin subunits α6 and αv. The prevalence of all these markers increased significantly in Myh11 lineage-positive pericytes (Figure 3 D, E, I, J). Representative plots of healthy and diseased lung for matrix-remodeling markers αSMA and Col1a1 (Figure 3B, C) demonstrate this shift. The amount of Myh11 lineage-positive pericytes positive for αSMA nearly doubles two weeks after bleomycin treatment (Figure 3D). Col1a1 is a collagen subunit that can be labeled intracellularly, provides a snapshot of cellular collagen synthesis, and is used as a measure of myofibroblastic tissue remodeling [44–47]. As with αSMA, the incidence of Col1a1+/Myh11 lineage-positive pericytes significantly increases by over two-fold in the bleomycin treatment group (Figure 3E). A tripling of the frequency of αSMA+/Col1a1+/Myh11 lineage-positive pericytes was observed (Figure 3F).

**Figure 3.**
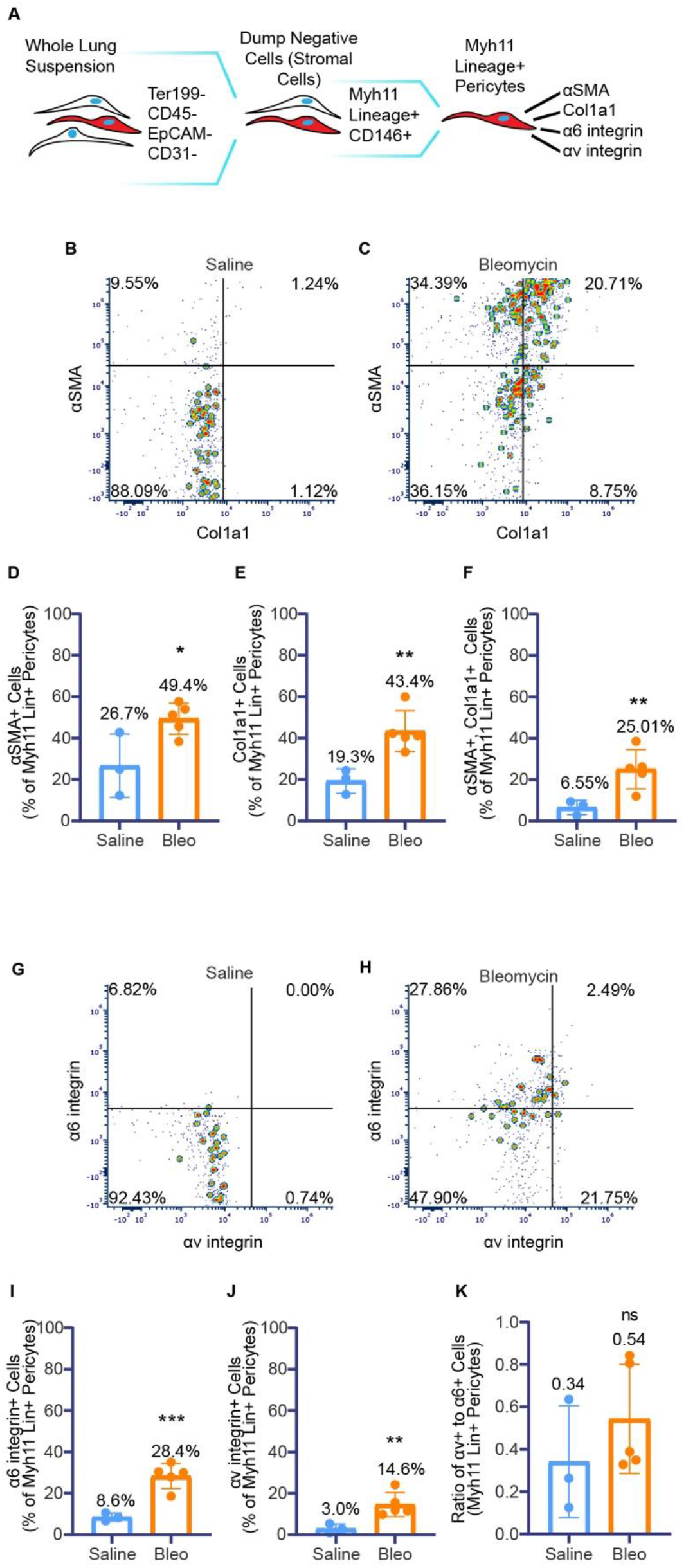
Flow cytometry of Myh11 lineage-positive pericytes isolated from saline and bleomycin treated mouse lungs show increased matrix-remodeling and matrix-adhesion proteins. (A) Gating hierarchy to isolate Myh11 lineage-positive pericytes for phenotyping. (B-F) Analysis of the expression of tissue-remodeling markers, αSMA and Col1a1, in Myh11 lineage-positive pericytes. Representative plots of αSMA and Col1a1 from saline (n=3, B) and bleomycin (n=5, C) treatment groups. Quantitation of αSMA-positive (D), Col1a1-positive (E), and double-positive (F) cells. (G-K) Analysis of adhesion integrins, α6 and αv, in Myh11 lineage-positive pericytes. Representative plots of α6 and αv from saline (G) and bleomycin (H) treatment groups. Quantitation of α6 integrin (I), αv integrin (J) and the ratio of αv positive cells to α6 positive cells (K). Data are expressed as means ± standard deviation. Statistical significance determined via unpaired, one-tailed student’s t-test. ns = not significant; p < 0.05 = *; p < 0.01 = **; p < 0.001 = ***.

Integrin subunits α6 and αv confer affinities for the basement membrane protein laminin and the RGD motif, respectively. Representative plots from saline and bleomycin treatments (Figure 3G, H) demonstrate significant population shifts between saline control and bleomycin-treated mice. The frequency of α6+/Myh11 lineage-positive pericytes increases threefold in the bleomycin treatment group (Figure 3I), and αv+/Myh11 lineage-positive pericytes increase nearly fivefold (Figure 3J). No significant change in the ratio of αv to α6 integrin-expressing, Myh11 lineage-positive pericytes between treatment groups can be seen (Figure 3K). This ratiometric quantification interrogates a shift in integrin expression across the population of Myh11 lineage-positive pericytes.

When Myh11 lineage-positive pericytes are compared to the broader population of stromal cells (defined as cells negative for lineages Ter119, CD45, CD31, EpCAM [44]), it can be observed that the relative ratio of αSMA+ pericytes to αSMA+ stromal cells decreases in bleomycin, even while the incidence of pericytes positive for αSMA increases (Supplemental Figure 2D). Myh11 lineage-positive pericytes comprise the bulk of Col1a1+ cells in the stromal population of both healthy and diseased lung (Supplemental Figure 2D).

### Myh11 lineage-positive pericytes isolated from fibrotic lungs are enriched for tissue-remodeling genes

Myh11 lineage-positive pericytes were isolated from whole-lung digestions that were obtained two weeks after bleomycin-treatment or saline-treatment, as described in Figure 4A. Cells were run through an Illumina sequencing platform, with details provided in the Methods section. A list of the top 30 ranked genes by fold change can be seen in Figure 4B, with an extended top 100 genes provided in Supplemental Figure 4A. The top of the list consists of several ECM components, matrix metalloproteinases (MMPs), and genes associated with activation of immune complement. A GeneSet Enrichment Analysis (GSEA) allows for an unbiased perusal of 1378 mapped mouse genesets. Our analysis found no significant enrichment of genesets in the saline treated-group, and 49 genesets were found to be enriched in the bleomycin-treated group. A visual aide to understanding the multiple tests and scores of bulk GSEA can be found in Supplemental Figure 4B. Genesets can be seen in Figure 4C and comprise Cellular Component (CC), Biological Processes (BP), Molecular Function (MF), and Matrisome (Matri) categories, of which the top five are shown. CC, MF, and BP are domains generated by the GeneOntology group [48,49], while Matri is a curated list of fibrosis and fibroproliferative-relevant genesets by Naba and Hynes [50]. Across all categories, significant enrichment of ECM and ECM-related processes is seen in cells procured from bleomycin-treated lungs. Also enriched are several cell cycle and proliferation genesets, implying a metabolically activated and mitotic cell population. The entire list of significant genesets can be found in Supplemental Figure 4C.

**Figure 4.**
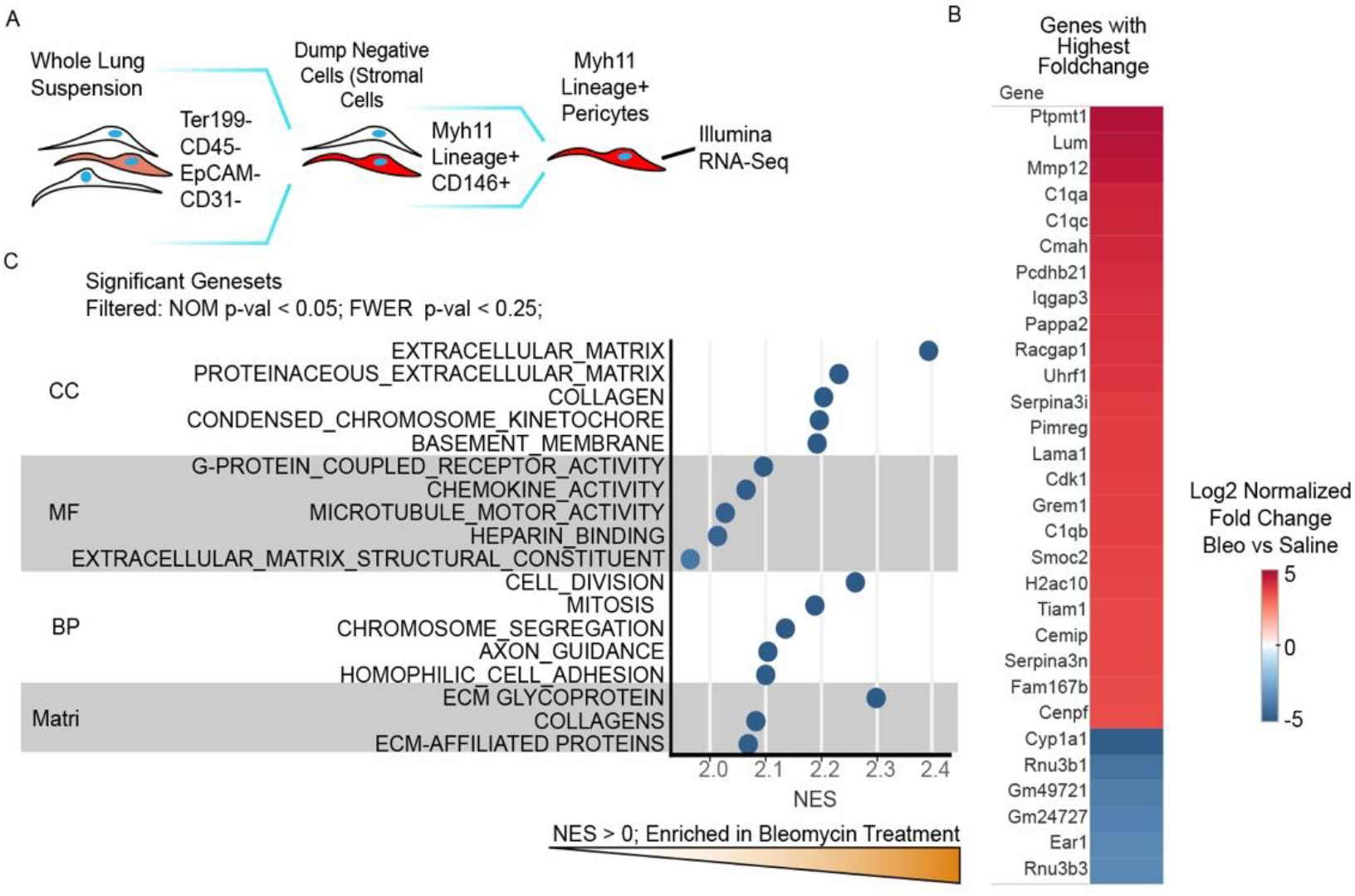
RNA sequencing (RNA-seq) of Myh11 lineage-positive pericytes isolated from saline and bleomycin treated mouse lungs demonstrate increased expression of tissue-remodeling genes in the disease model. (A) Gating hierarchy to isolate Myh11 lineage-positive pericytes for phenotyping. (B) Heat map of top 30 most differentially expressed genes out of 22,203 between pericytes from bleomycin (Bleo) and saline groups. (C) GeneSet Enrichment Analysis (GSEA) of RNA-seq data in Myh11 lineage-positive pericytes. The threshold for significance of tested genesets is nominal p-value (NOM p-val) < 0.05 and Family-wise Error Rate (FWER) < 0.25. There is no significance threshold for Normalized Enrichment Score (NES), as it is a measure of GO set expression on phenotype. The top five genesets from each domain meeting the filter criteria are displayed. All shown genesets have a NES score > 0, indicating all significant GO sets are enriched in bleomycin treatment. Cellular component (CC), molecular function (MF), biological process (BP), and matrisome denote separate categorical domains of gene ontologies.

### Primary Myh11 cell culture on fibronectin with RGD-inhibition reveals RGD-dependent increase of cellular αSMA

Fibronectin coated, stiff substrates are thought to activate myofibroblasts through αv integrin focal adhesions [51]. A laminin coating with no readily available RGD integrin ligand was chosen for a negative control, as pericyte α6 investment in the laminin-rich basement membrane is a known requirement for cellular and tissue homeostasis and does not activate myofibroblastic phenotypes [33,52]. Cyclic RGD (cRGD) in an approximately 100-fold molar excess beyond reported IC50 values for αv heterodimer adhesion was used to prevent RGD engagement, with nonbinding cyclic RAD (cRAD) used as a control [53]. Mouse fibronectin or laminin-coated cover slips seeded with CD146+ MACS-enriched primary cells isolated from digested lung can be seen after 24 hours culture (Figure 5A). The prevention of cellular engagement with RGD results in less spread and less contractile cells, as observed by αSMA. The mean fluorescence intensity (MFI) of cellular αSMA across cell preparations from several mice (n=3) can be seen in Figure 5B. The Myh11 lineage-positive cells plated on fibronectin with cRAD nonblocking control generated significantly increased αSMA compared to the cRGD blocked group (Figure 5B). The laminin surface controls show no αSMA increases relative to the fibronectin surface with either treatment and are shown together as “LAM + Peptide”.

**Figure 5.**
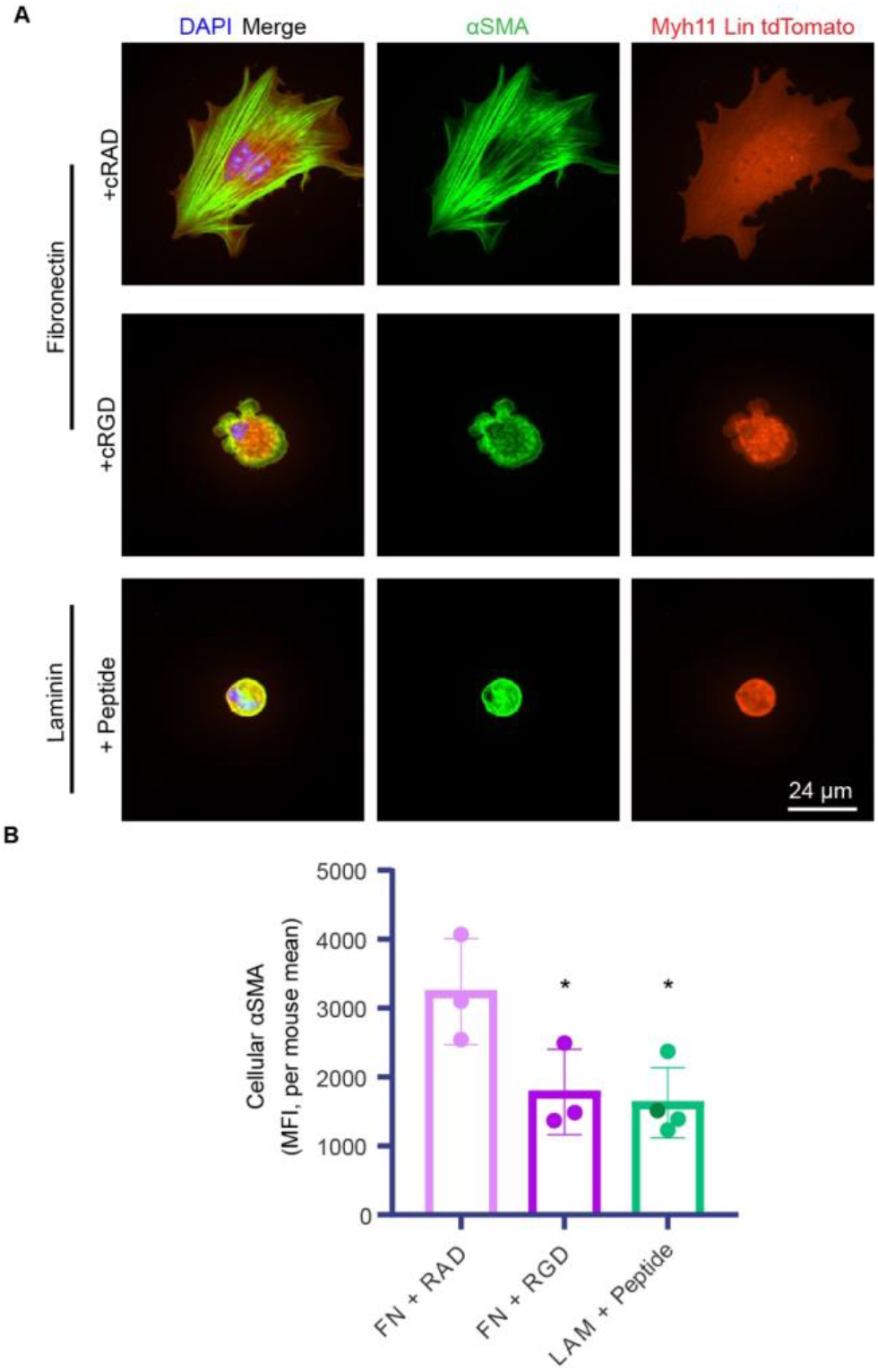
Cell culture of Myh11 lineage-positive cells with RGD inhibition reveals RGD-dependent increase of αSMA on fibronectin-coated substrates but not on laminin-coated substrates. (A) Representative immunofluorescence (IF) micrographs of Myh11 lineage-positive cells (red, endogenous fluorophore) and αSMA (green). Cells were treated with the soluble RGD inhibitor, cyclo-RGDfK (cRGD), or the noninhibitory control cyclo-RADfK (cRAD). (B) Quantitation of mean fluorescence intensity (MFI) of αSMA within each cell. Data are expressed as means of cells from individual mice (n=3), with 16-60 cells averaged per mouse, ± standard deviation. No differences were seen between peptides in the laminin group, and they were thus merged. Dark green = LAM + RGD; light green = LAM + RAD. Statistical significance was determined via unpaired, one-tailed student’s t-test. ns = not significant; p < 0.05 = *; p < 0.01 = **; p < 0.001 = ***.

## Discussion

We demonstrate the ability to identify pericyte-derived myofibroblasts using a pericyte lineage reporter mouse treated with bleomycin to induce lung injury. Using this model system, we have identified the pulmonary pericyte as a population of myofibroblast progenitors. We showed that pericytes adopt a tissue-remodeling phenotype in bleomycin-induced lung injury, and we identified the extracellular matrix ligand (RGD) in fibronectin as a potentiator of pericyte-to-myofibroblast transition *in vitro*.

The body of work describing pericyte contributions to fibrosis, and the pericyte-to-myofibroblast transition, is still nascent [18,20,22,54–57]. Foundational studies have shown pericytes can transition into myofibroblasts in disease [20,54,56], with additional research exploring possible molecular mechanisms driving the transition [22] and pericyte-specific knockdown studies, which confer protection in fibrotic disease models [37,41,57]. The detailed characterization of pericytes in this work through histological analysis, high-dimensional flow cytometry, and gene ontology RNASeq analyses provides an unprecedented level of insight into the response of pericytes in the early phases of repair following acute lung injury. Our characterization of pericytes using a variety of surface markers through extensive cytometric phenotyping contributes to the existing literature by comparing them to the broader stromal cell population. The ability to contextualize our population of interest among the broader stromal cell population allows for relative comparisons to be made between our pericyte lineage and the broader stromal cell population, something not possible without high-dimensional cytometric analyses. The abundance of pericyte collagen production (Col1a1+) within the larger stromal population agrees with prior pericyte research in kidney fibrosis [47]. Contrastingly, pericytes make up a much smaller proportion of the stromal contractile (αSMA+) population. This heterogeneity in relative contributions of myofibroblast markers to stromal cell population provides further evidence to support the growing understanding of fibroblast heterogeneity, which has mostly been obtained via single-cell RNA sequencing, showing that myofibroblasts originate from a variety of cell populations. This new focus on myofibroblastic heterogeneity is revealing that phenotypes once considered pan-myofibroblastic (based on whole population analyses) can be attributed to distinct subpopulations of myofibroblasts [45,58,59]. Our data suggest that pericytes are the primary collagen 1-producing stromal cells in both healthy and bleomycin-treated lung.

Our work examines integrin expression and activation for the first time in the murine Myh11 lineage model of pericytes. Integrins are known to be important in mediating fibroblast differentiation into myofibroblast phenotypes [24,37,51,60], so we posited that they might play a role in the pericyte-to-myofibroblast transition. Our findings show a pronounced increase in pericyte αv and α6 protein expression. This enhanced adhesion profile is not unexpected; ligand density for ECM-binding integrins increases in fibroproliferative injury [61,62]. While the ratio of αv to α6 surface expression is not significantly altered in the pulmonary pericyte population of murine lungs treated with bleomycin, the amount and activation/engagement of αv(β3) are both significantly increased in the injury model. The lack of significant increase in total fibronectin proximal to pericytes following bleomycin induced injury is curious, especially in light of significantly heightened levels of active αvβ3 integrin on the surface of pericytes. However, the bleomycin model of lung injury is a resolving model and these data may point to a more balanced fibroproliferative repair compared to pathological, chronic remodeling observed in human fibrotic diseases. These data further suggest additional regulatory mechanisms are at play that contribute to αvβ3 activation in the pericyte population, including known inside-out mechanisms of integrin activation due to exogenous agonists that are known to be present in the provisional wound repair environment, such as thrombin and others [53,63]. We might also speculate, based on our previous work, that contractile (αSMA+) pericytes engage a known integrin switch driven through a mechanically sensitive conformational change within fibronectin’s integrin binding domain that drives a strong preference for fibronectin-αvβ3 engagement [24,25,51]. In total, these data strongly suggest that there is a mechanically responsive/active, provisional matrix-engaged pericyte population during lung injury repair.

Studies of pericyte behaviors have been historically limited by the available approaches for identifying and tracking them in living tissues. Given the wide range of non-unique surface markers that pericytes express [14], morphologic criteria and physical orientation relative to capillary endothelium have been used to positively identify pericytes [42]. However, this morphologic description is limiting when the goal is to interrogate pericyte transitions into other cell types. The Myh11 lineage system is one of many published murine models [16] that has provided a means to identify this cell population and its lineage *in vivo*. However, we cannot assume that the pericytes labeled by this Myh11 lineage tracing system are identical to those labeled by other lineage tracing systems, nor can the Myh11 lineage be assumed to universally label all pericytes. Since the Myh11 lineage cell population also includes smooth muscle cells (SMCs) [21,64,65], and since SMCs have been shown to also express CD146 [66], it is possible that our RNAseq and cytometric results include contributions from SMCs, in addition to pericytes. Given this caveat, we can still conclude that the mural cell population, which includes both pericytes and SMCs, are active participants in fibrotic remodeling following bleomycin-induced injury in the lung. And, considering our RNAseq and cytometric data in light of our histological data, which demonstrate based on cell morphologies and proximity to capillaries that pericytes are the predominant cells exhibiting myofibroblastic behaviors, we can conclude that if SMCs also undergo myofibroblast differentiation, their role in fibrotic tissue remodeling is relatively minor compared to that of pericytes.

We explored αv integrin, a fibronectin/provisional matrix binding protein known to potentiate myofibroblast phenotypes, in the context of pericyte-to-myofibroblast differentiation. However, αv does not only bind to RGD, but has many ligands, and studies of αv in the microcirculation tend to focus on αv-platelet endothelial cell adhesion molecule (PECAM) interactions. We do not evaluate the contributions of endothelial cell remodeling or behaviors to pericyte-to-myofibroblast transitions, but we acknowledge the remodeling of microvessel networks as essential components of lung fibrosis [67].

We acknowledge there are limitations in our model systems and techniques which must be considered in the interpretation of our results. The bleomycin injury model for resolving lung fibrosis has been extensively used and cited in literature, but does not accurately recapitulate many aspects of human lung fibrosis [43,68,69]. It should be thought of as a platform to study cell and tissue-scale phenomena in fibroproliferative injury, which has enabled our extensive research on myofibroblasts in fibrotic lung injury [26,27,51]. The cell dissociation process necessary for preparing cells for cytometry sorting and analyses, including RNASeq, subjects the isolated cells to a brief period of enzymatic and mechanical perturbation, which may skew data in unforeseen ways, though care was taken to limit known artifactual stimulants of cells, such as titration of digestion and lysis steps, and addition of DNAse and EDTA to sorting and digestion steps. Additionally, published research verifies the stability of many of our surface markers in our digestion model [70]. The spectral flow cytometry we employed provides an unprecedented, highly dimensional cytometric dataset, but potential fluorescence overlap limits the number of cell populations that can be accurately compared within a given gating hierarchy.

Translating known downstream signaling of integrins in fibroblasts to other stromal cell populations, including pericytes, has provided some insight into biologic processes required for the generation of myofibroblastic cell phenotypes from non-fibroblast precursor populations. Myocardin-related transcription factor (MRTF) is an essential component of integrin-mediated myofibroblast activation, and recent studies have shown its requirement for various stromal cell to myofibroblast transitions [22,71]. Likewise, YAP/TAZ/Hippo signaling is known to regulate myofibroblast phenotype and is dysregulated in disease [72], while studies of YAP/TAZ deficient pericytes have demonstrated the loss of YAP/TAZ in Gli1 perivascular stromal cells is protective in fibrotic injury of kidneys [73,74]. The increase in active pericyte αvβ3 measured in this study is known to potentiate those signaling cascades, but direct measurement of these pathways was outside the scope of this manuscript.

Many other opportunities to investigate myofibroblast biology as it pertains to pericytes are enabled by the Myh11 pericyte lineage, and this will allow for a more complete understanding of the process by which pericytes and other stromal cells can be driven towards pathologic myofibroblastic behavior. While myofibroblast responses to various collagenous and fibronectin-rich substrates are well understood [25,62,75], pericyte responses to the changes in ECM composition during injury are less understood. The study of basement membrane remodeling in fibrotic injury as it coincides with generation of provisional matrix has begun here with our evaluation of fibronectin in locations that are proximal to pericytes. Indeed, our findings link ECM-based integrin ligand RGD with cell surface αv integrin activation in the pericyte-to-myofibroblast transition. Further study into the mechanisms underpinning phenotypic changes in pericytes during lung fibrosis may reveal new potential diagnostic and therapeutic targets for lung fibrosis.

## Materials and Methods

A full materials and methods, including antibodies and dilutions, can be found in the supplement.

### Mice

*All procedures were performed in accordance with the Institutional Animal Care and Use Committee of the University of Virginia*.

### Induction of Lineage

*Myh11-CreERT2 ROSA STOPfl/fl tdTomato* mice 6-12 weeks of age were injected intraperitoneally with 1mg tamoxifen each day for 10 days over two weeks.

### Tracheal Bleomycin Administration

Animals were anesthetized with a ketamine/xylazine cocktail. Animals were hung by their incisors at 45 degrees. Bleomycin sulfate (1-3 U/kg) (Meitheal Pharmaceuticals, Chicago, IL, USA) in saline or sterile saline (2uL/gm) was instilled into the lungs through the trachea through angiocatheter tubing placed down the animal’s throat.

### Harvest of Lung Tissuse

Animals were euthanized at two weeks post administration of bleomycin. The thorax was opened and lungs perfused with sterile PBS through the heart until cleared of blood. Submandibular tracheotomy was performed and the lungs lavaged with PBS to clear airway cells. Lungs were inflated with 2% agarose for histology or placed into PBS for dissociation.

### Histologic Staining

Lungs for histologic analyses were submerged in 4% paraformaldehyde (PFA) for 30 minutes or were fresh frozen. Lungs were sectioned, and sections are blocked and stained as appropriate for the experiment and finally hard mounted for imaging.

### Immunofluorescence Imaging

Images captured on an UltraView Vox Spinning Disk Confocal Microscope (PerkinElmer, Waltham MA, USA) using Volocity 6.3.1 software (PerkinElmer) and Nikon PlanFluor 20x and Nikon Apo TIRF 60x objectives, or a BZ-X810 widefield fluorescence microscope (Keyence, Osaka, Japan). In-house software (https://bitbucket.org/pythoncardiacmodel/publicpythoncardiacmodel/src/master/) and Volocity used for image stitching. Analyses and particle counts performed in Volocity or Fiji/Imagej [76].

### Lung Dissociation/Single Cell Isolation

Lung lobes/large chunks from a single mouse were placed into into a 2ml microfuge tube and chopped into small (< 2mm) chunks with sterile scissors. 1ml of digestion solution (TM Liberase at 4 units/mL (Roche, Basel, Switzerland) and DNAse Type I at 800 units/mL (ThermoFisher) made in sterile PBS) was added to each tube. Tubes were placed on a rotisserie at 37C for 20 minutes. Digest was then mechanically ground through 100um nylon filters (ThermoFisher) and placed in ACK RBC Lysis buffer (ThermoFisher) for 3 minutes at RT or as needed. Cells were pelleted and, if further cleanup is needed, a densisty gradient was used (debris removal soluton, Miltenyi, Bergisch Gladbach, GER) according to manufacturer direction.

### Cell Culture

Primary cell culture performed in DMEM (ThermoFisher, Waltham MA, USA) with fibronectin-depleted (depleted via column purification using gelatin sepharose) FBS at 10% on glass coverslips coated with 10ug/mL murine fibronectin (native mouse fibronectin, Abcam, Cambridge MA, USA) or laminin (laminin mouse protein, natural, ThermoFisher). Inhibition studies included 100 μM of Cyclo(-RGDfK) or Cyclo(-RADfK-) (Anaspec, Fremont CA, USA) for the incubation. Cells fixed at 24 hours.

### Flow Cytometric Analyses

Live cell sorting performed on Miltenyi AutoMACS and BD Influx sorters. CD146 (LSEC) MACS Beads (Miltenyi) added to cells according to manufacturer directions and sorted in a poseld2-protocol on the AutoMACS.

Phenotyping panels performed on Cytek Aurora spectral flow cytometers with fixing and permeablizing by Fix&Perm kit (ThermoFisher).

### RNA Sequencing and Analysis

RNA Sequencing and Analysis is described in the supplement.

## Supporting information

Manuscript Supplement

## Acknowledgements

University of Virginia Flow Cytometry Core Facility (FCCF).

Chris Waters for the use and instruction of image stitching application (https://bitbucket.org/pythoncardiacmodel/publicpythoncardiacmodel/src/master/).

Sue Landes, Chiuan-Ren (Vincent) Yeh, and Anthony Bruce for their assistance in animal studies and mouse colony maintenance.

The entirety of Peirce-Cottler and Barker labs for their assistance in hypothesis generation and troubleshooting.

## Funding

The Hartwell Foundation and U01AR069393 (SMP); R01HL127283 and R01HL132585 (THB); UVA Fibrosis Initiative

## Author Contributions

RTH designed and performed experiments and analysis, and wrote the manuscript AEM assisted with RNASeq processing and analysis and manuscript RH and CS assisted in experimentation and image analysis THB and SMP provided editorial support and assisted with experimental design

## Abbreviations

ECM: Extracellular Matrix
αSMA: Alpha smooth muscle actin
SMC: Smooth muscle cell

## Notes

### Competing Interest Statement

The authors have declared no competing interest.

