## Supplementary material for "Extracellular Matrix Remodeling Associated with Bleomycin-Induced Lung Injury Supports Pericyte-To-Myofibroblast Transition": Manuscript Supplement

SUPPLEMENTAL

**Supplemental Figure 1. The Myh11-CreERT2 ROSA STOPfl/fl tdTomato reporter mouse requires tamoxifen for reporter induction and does not alter lung histology.** (A) Schematic of the Myh11-CreERT2 ROSA STOPfl/fl tdTomato lineage reporter mouse (B) Representative immunofluorescence (IF) and hematoxylin and eosin (H&E) micrographs of lung. IF histology stained for tdTomato (red, endogenous fluorophore), and DAPI (blue).

**Supplemental Figure 2. Broader phenotyping of myh11 lineage pericytes and lineage pericytes in relation to the putative stromal cell population as identified by flow cytometry.**

(A) Gating hierarchy to identify Myh11 lineage-positive pericytes for phenotyping. (B) Additional surface markers assayed (C) Gating hierarchy to isolate putative stromal cells via negative marker selection. (D) Comparison of saline and bleomycin marker profiles on both stromal cells and the perivascular subset of that same population.

**Supplemental Figure 3. Fluorescence Minus Ones (FMOs) and positive gating for flow cytometric probes used in Figure 3.** Representative plots of the gating strategy for cellular phenotyping. Some positive gates were simplified to rectangles for readability.

**Supplemental Figure 4**. **RNA sequencing (RNA-seq) of myh11 lineage pericytes isolated from saline and bleomycin treated mouse lungs demonstrate increased expression of tissue-remodeling genes in the disease model.** (B) Heat map of top 100 most differentially expressed genes out of 22,203 between pericytes from bleomycin (Bleo) and saline treatment groups. (C) GeneSet Enrichment Analysis (GSEA) plot of 1378 Gene Ontology (GO) sets, visualizing the test statistics: nominal p-value (NOM p-val); Normalized Enrichment Score (NES); Family-wise Error Rate (FWER). The threshold for significance is highlighted green (NOM p-val < 0.05) and white-to-blue color dots (FWER < 0.25). There is no signficance threshold for NES, as it is a measure of GO set expression on phenotype. (D) GO sets meeting the filter criteria (NOM p-val < 0.05; FWER < 0.25) are displayed. All sets have a NES score > 0, indicating all significant GO sets are enriched in bleomycin treatment. Cellular component (CC), molecular function (MF), biological process (BP), and matrisome denote separate categorical domains of gene ontologies.

**Supplemental Table 1: Antibodies**

| **Antibody** | **Clone** | **Dilution** | **Product Info** |
| --- | --- | --- | --- |
| **Flow** | | | |
| aSMA | 1A4 | 1/200 | NBP2-34522 |
| a6 integrn/CD49f | GoH3 | 1/100 | Bio legend 313612 |
| av integrin/CD51 | RMV7 | 1/100 | BD 551380 |
| PDGFRB/CD140b | APB5 | 1/100 | Bio Legend 136008 |
| PDGFRA/CD140a | APA5 | 1/100 | ThermoFisher 25-1401-82 |
| CD31 | MEC 13.3 | 1/50 | BD 612802 |
| EPCAM/CD326 | EBA-1 | 1/100* | BD 743544 |
| CD146 | ME-9F1 | 1/100 | BD 562232 |
| CD45 | 30-F11 | 1/100* | Bio Legend 103147 |
| Ter119/RBC | TER-119 | 1/100* | BD 740686 |
| Col1a1 | poly | 1/100 | ABNova PAB17204 |
| L/D NIR | n/a | as directed | ThermoFisher L34975 |
| *** combined dilution of all 'dump' Abs is 1/100, each individual Ab is <1/100** | | | |
| **Histology** | | | |
| CD31 | MEC 13.3 | 1:100 | BioLegend 102504 |
| aSMA | 1a4 | 1:200 | NBP2-34522 |
| fibronectin | poly | 1:200 | Abcam ab2413 |
| active avb3 integrin/Wow1 | n/a | 1:200 | gift of Sanford Shattil, University of California, San Diego |
| **secondaries** | | 1:1000 | AF conjugates |

Supplemental Figures

Supplemental Figure 1

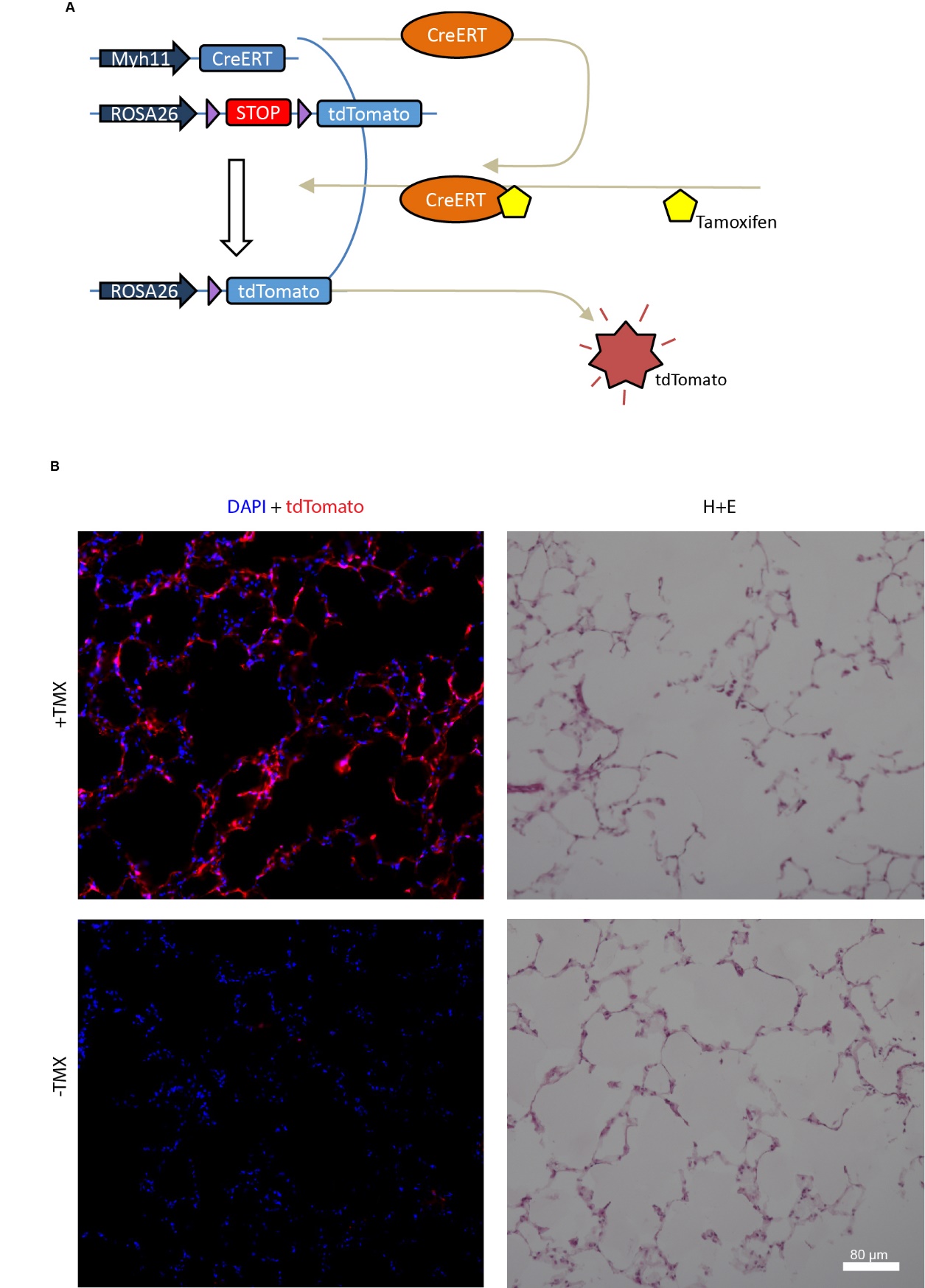

Supplemental Figure 2

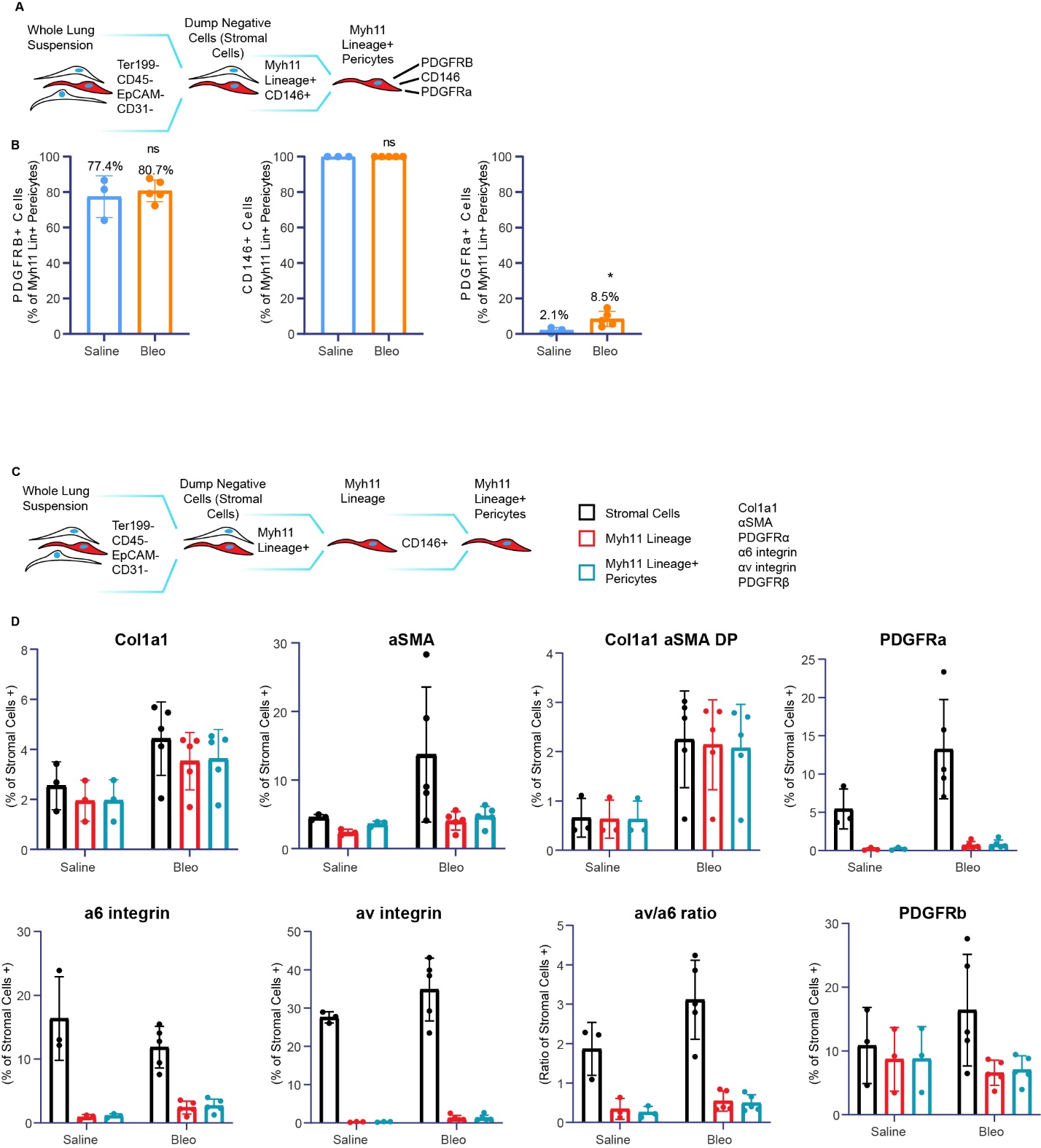

Supplemental Figure 3

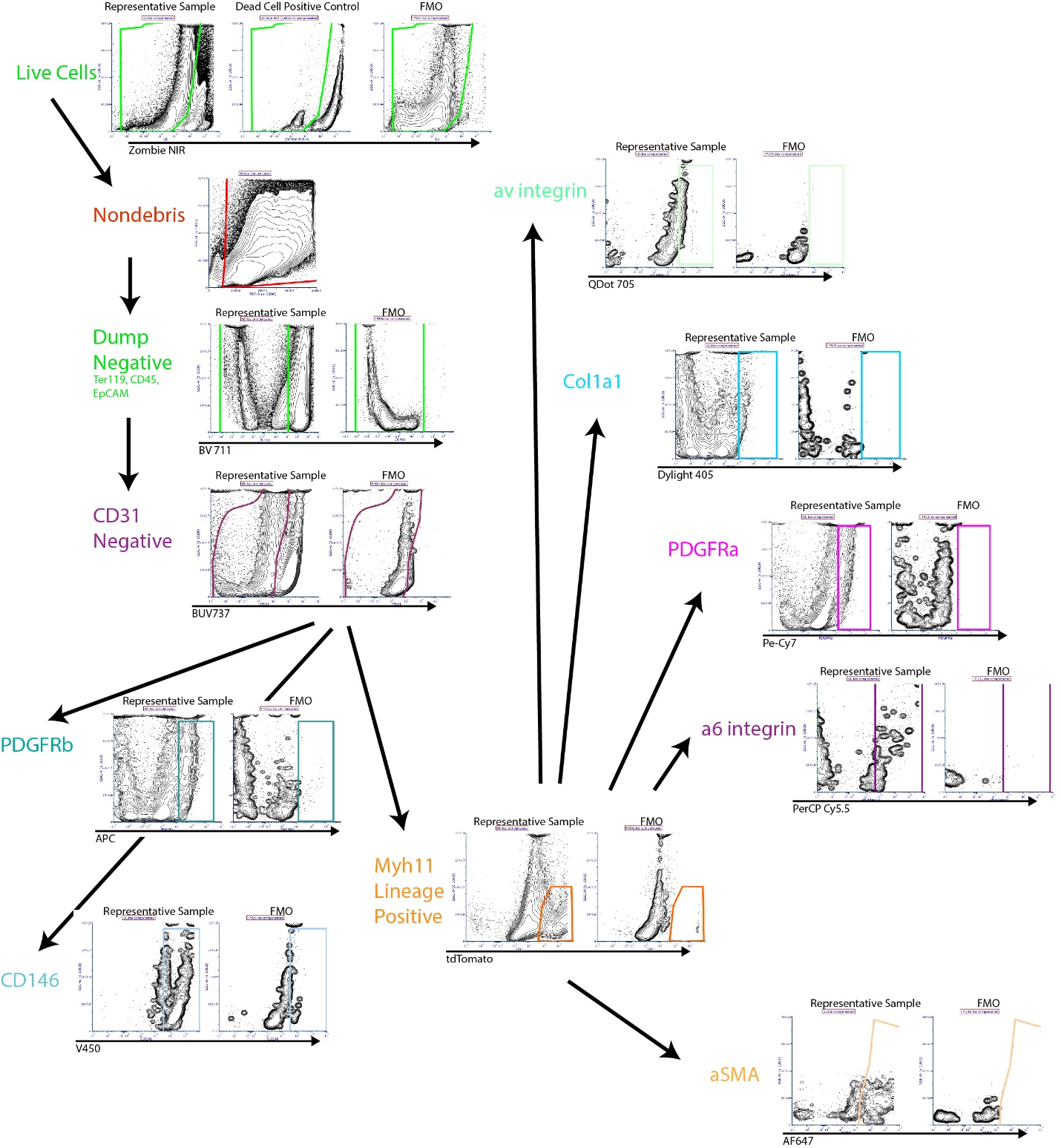
Supplemental Figure 4

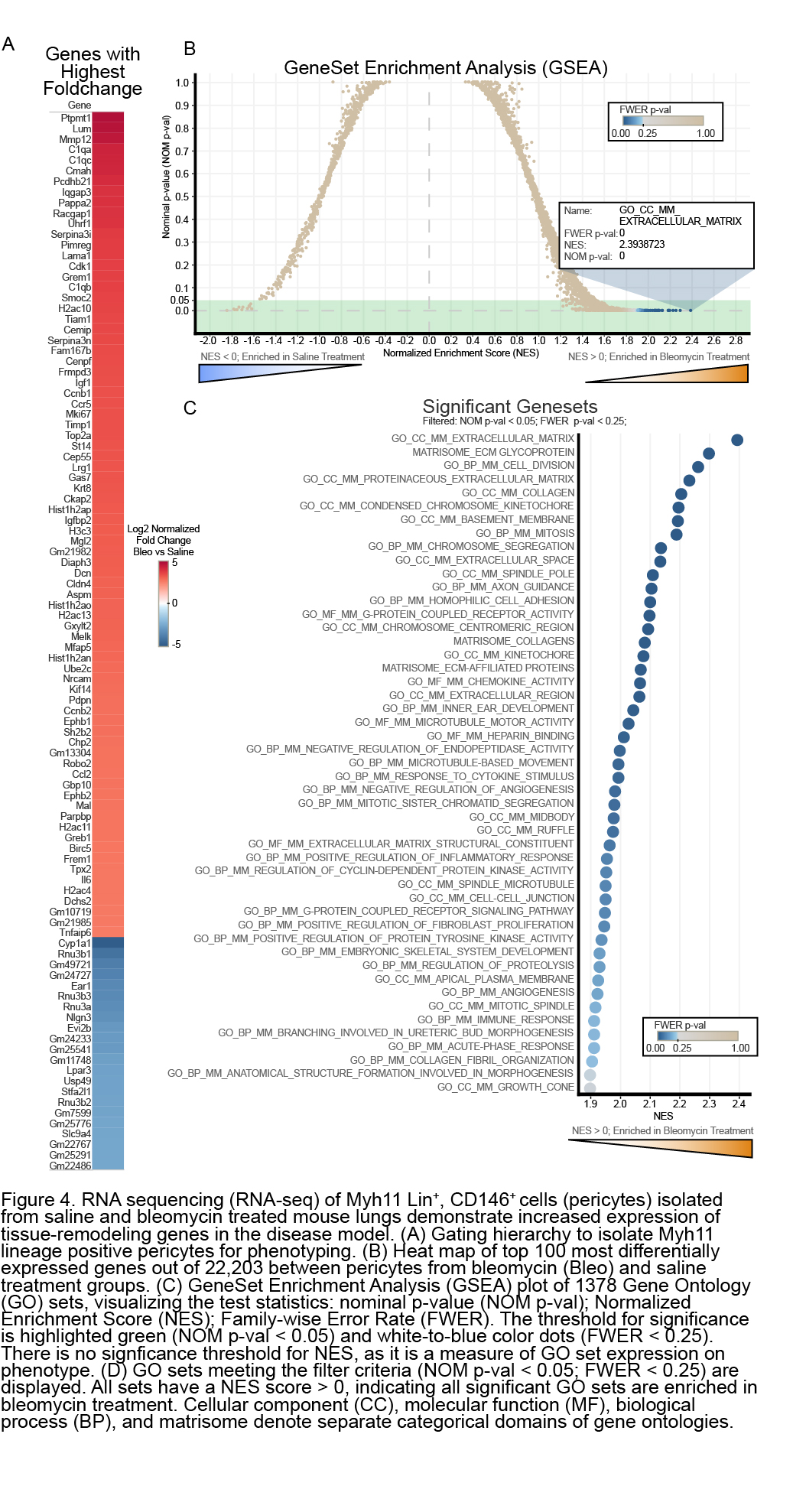

**Materials and Methods Supplement**

*Mice*

*All procedures were performed in accordance with the Institutional Animal Care and Use Committee of the University of Virginia. Myh11-CreERT2 ROSA STOPfl/fl tdTomato* mice 6-12 weeks of age were injected intraperitoneally with 1mg tamoxifen (10mg/mL of tamoxifen (Sigma-Aldritch, St.Louis, MO, USA) in 100 uL peanut oil (Sigma-Aldritch) each day for 10 days over two weeks.) Mice were rested for four weeks to allow clearance of tamoxifen, oil, and transient circulating cells before subsequent procedures.

*Tracheal Bleomycin Administration*

Animals were anesthetized with a ketamine/xylazine cocktail (60-80/5-10 mg/kg). Animals were placed on a commercial board from Hallowell EMC and hung by their incisors at 45 degrees. Bleomycin sulfate (1-3 U/kg) (Meitheal Pharmaceuticals, Chicago, IL, USA) in saline or saline vehicle control (2uL/gm) was instilled into the lungs through the trachea through angiocatheter tubing placed down the animal’s throat and connected to a 1mL syringe. Mice were monitored during and post- procedure to ensure recovery from anesthesia and not returned to housing until they were fully ambulatory and breathing normally.

*Harvest of Lung Tissue*

Animals were euthanized two weeks post-bleomycin via CO2 asphyxiation with secondary cervical dislocation. Mice were sprayed with 70% ethanol to sterilize and mat fur, and an incision was made through the skin extending from 5mm below the sternum to the mandible. A bilateral incision was made perpendicular from the abdominal end of the previous incision, exposing the caudal edge of the ribcage and the abdominal cavity and peritoneum. The peritoneum along the caudal ribcage edge was cut along the entire length of the ribcage. The diaphragm was dissected away and the ribcage cut or moved away to expose the cardiopulmonary unit. The inferior vena cava was cut and the heart perfused with 5-10ml of sterile saline or PBS, until blanching of the lungs was complete.

In the submandibular area, salivary glands were dissected away and the trachea cannulated with a blunt tipped needle or trachea tube and secured with suture. Lungs were then washed 3x with sterile saline or PBS, and then inflated with 2% UltraPure LMP Agarose (ThermoFisher, Waltham, MA, USA) for histology or sterile saline/PBS for lung dissociation. The entire cardiopulmonary unit was dissected out of the thorax and placed into saline/PBS on ice for subsequent protocols.

*Histologic Staining*

Lungs for histologic analyses were submerged in 4% paraformaldehyde (PFA) for 30 minutes or were fresh frozen. Lungs were placed in 30% sucrose solution until they sink, at which point they were embedded in OCT and frozen. Lungs were cryosectioned at 8 or 30 micron thickness for histochemical and immunofluorescent staining.

Slides for histochemical analyses were and stained with hematoxylin and eosin (Ricca, Arlington TX, USA) and Picrosirius Red (Abcam, Cambridge MA, USA) following manufacturer protocols.

All steps performed in a hydration chamber. All antibodies and dilutions are listed in Table S1. OCT was washed away with PBS and section was fixed (if necessary) with 4% PFA or methanol. Section was then permeabilized for 1 hour with 0.5% TritonX-100 in PBS. Section was then blocked with the mouse serum (if not available, FBS) at 5% and the secondary antibody host serum at 5% in PBS for 1 hour at RT or overnight at 4C. Primary antibodies added for 3 hours RT or overnight at 4C. Section was washed with PBS for 5m 5x. Secondaries and conjugated primaries were added for 3 hours RT or overnight at 4C. Section was washed with PBS for 10m 5x. Tissue was washed and mounted with Prolong Diamond (ThermoFisher). DAPI Counterstain from mounting with Prolong Diamond with DAPI (ThermoFisher)

*Immunofluorescence Imaging*

Images captured on an UltraView Vox Spinning Disk Confocal Microscope (PerkinElmer, Waltham MA, USA) using Volocity 6.3.1 software (PerkinElmer) and Nikon PlanFluor 20x and Nikon Apo TIRF 60x objectives, or a BZ-X810 widefield fluorescence microscope (Keyence, Osaka, Japan) Tiled images taken at 10% overlap and stitched using in-house software (<https://bitbucket.org/pythoncardiacmodel/publicpythoncardiacmodel/src/master/>) and Volocity. Analyses and particle counts performed in Volocity or Fiji/Imagej [76].

*Lung Dissociation/Single Cell Isolation*

All steps performed on ice unless otherwise indicated. Heart, fat, connective tissue and primary bronchi were dissected away from the lung lobes, minimizing non-lobe tissue. Lung lobes/large chunks from a single mouse were placed into into a 2ml microfuge tube and chopped into small (< 2mm) chunks with sterile scissors. 1ml of digestion solution (TM Liberase at 4 units/mL (Roche, Basel, Switzerland) and DNAse Type I at 800 units/mL (ThermoFisher) made in sterile PBS) was added to each tube. Tubes were placed on a rotisserie at 37C for 20 minutes. Digest was then mechanically ground through 100um nylon filters (ThermoFisher) and placed in ACK RBC Lysis buffer (ThermoFisher) for 3 minutes at RT or as needed. Cells were pelleted and, if further cleanup was needed, a densisty gradient was used (debris removal soluton, Miltenyi, Bergisch Gladbach, GER) according to manufacturer direction.

*Cell Culture*

Primary cell culture performed in DMEM (ThermoFisher, Waltham MA, USA) with fibronectin-depleted (depleted via column purification using gelatin sepharose) FBS at 10% on glass coverslips coated with 10ug/mL murine fibronectin (natural mouse fibronectin, Abcam) or laminin (laminin mouse protein, natural, ThermoFisher). Inhibition studies included 100 µM of Cyclo(-RGDfK) or Cyclo(-RADfK-) (Anaspec, Fremont CA, USA) in PBS to single cell suspensions immediately prior to plating. Cells allowed to adhere overnight, and media was replaced. Cells fixed at 24 hours, permeablized, and stained.

*Flow Cytometric Analyses*

All antibodies and dilutions are listed in Table S1.

Live cell sorting performed on AutoMACS (Miltenyi) and BD Influx (BD Biosciences, San Jose, CA, USA) sorters. Cells suspended in FACS buffer (PBS+5% FBS+1mM EDTA) for staining and sorting. CD146 (LSEC) MACS Beads (Miltenyi) added to cells according to manufacturer directions and sorted in a poseld2-protocol on the AutoMACS. Gating strategies for Influx can be seen in figure S6

Phenotyping panels performed on Cytek Aurora spectral flow cytometers with fixing and permeablizing by Fix&Perm kit (ThermoFisher). Cells suspended in FACS buffer (PBS+5% FBS+1mM EDTA) for staining and sorting.

RNA Sequencing

Cells isolated from murine lung were used to generate mRNA libraries for RNA sequencing. Extraction and library prep were performed commercially with rRNA depletion (GeneWiz, South Plainfield, NJ, USA). All libraries were sequenced using single lane of the Illumina HiSeq sequencer to generate 150bp paired-end reads at a total read depth of 350 million reads (GeneWiz). Following sequencing, the resulting fastq files were processed to remove low-quality reads (Phred quality score < 20) and the presence of any adapter sequences using TrimGalore [1]. The resulting quality of each sample was independently evaluated using FastQC [2]. Files that passed QC were further processed to obtain gene expression counts following a previously defined protocol by Pertea et al. [3]. Briefly, reads were aligned to the mouse genome (UCSC mm10) using the HISAT2 aligner [4] and transcripts assembled with StringTie [5]. The resulting StringTie output was used to produce read coverage tables for input into DESeq2 [6].

RNA Data Analysis

DESeq2’s median of ratios method was used to normalize for differences in sequencing depth and RNA composition between samples. Normalized counts were subsequently used to quantify differential expression in the diseased state (bleomycin vs. saline control). All gene ontologies were curated from The Gene Ontology (GO) Consortium or the MSigDB NABA_MATRISOME gene set [9], with murine GO homologes sourced from Xijin Ge [10]. Gene set enrichment analysis (GSEA) was used to identify significantly enriched gene sets performed using the GSEA-MSigDB desktop application and default parameters.

*Statistical analyses*

A two-way ANOVA or a Student’s t-test was performed using Prism (GraphPad, San Diego CA, USA), as indicated in each figure caption. Statistical significance was asserted at p-values < 0.05. All data are presented as average + standard deviation.

1. Krueger, Felix. Trim galore. A wrapper tool around Cutadapt and FastQC to consistently apply quality and adapter trimming to FastQ files 516 (2015): 517.
2. Andrews, Simon. FastQC: a quality control tool for high throughput sequence data. (2010).
3. Pertea, M., Kim, D., Pertea, G. *et al.* Transcript-level expression analysis of RNA-seq experiments with HISAT, StringTie and Ballgown. *Nat Protoc* **11,** 1650–1667 (2016). <https://doi.org/10.1038/nprot.2016.095>
4. Kim, Daehwan, Ben Langmead, and Steven L. Salzberg. HISAT: a fast spliced aligner with low memory requirements. *Nature methods* **12**(4), 357-360 (2015).
5. Pertea, M., Petrea, GM., Antonescu, CM. *et al.* StringTie enables improved reconstruction of a transcriptome from RNA-seq reads. *Nature biotechnology* **33**(3), 290 (2015).
6. Love, M. I., Huber, W., & Anders, S. (2014). Moderated estimation of fold change and dispersion for RNA-seq data with DESeq2. *Genome biology*, *15*(12), 550.
7. Naba A, Clauser KR, Hoersch S, Liu H, Carr SA, Hynes RO. The matrisome: in silico definition and in vivo characterization by proteomics of normal and tumor extracellular matrices. *Mol Cell Proteomics*. 2012;11(4):M111.014647. doi:10.1074/mcp.M111.014647
8. Subramanian, Aravind, et al. "Gene set enrichment analysis: a knowledge-based approach for interpreting genome-wide expression profiles." *Proceedings of the National Academy of Sciences* 102.43 (2005): 15545-15550.
9. Mootha, V., Lindgren, C., Eriksson, K. *et al.* PGC-1α-responsive genes involved in oxidative phosphorylation are coordinately downregulated in human diabetes. *Nat Genet* **34,** 267–273 (2003).
10. <http://ge-lab.org/#/data>
